# Taxonomic description and genome sequence of *Christensenella intestinihominis* sp. nov., a novel cholesterol-lowering bacterium isolated from human gut

**DOI:** 10.1101/2020.11.14.383166

**Authors:** Yuanqiang Zou, Wenbin Xue, Xiaoqian Lin, Guangwen Luo, Mei Lv, Ying Dai, Liang Xiao

**Author notes:** **Corresponding authors**: Liang Xiao., Address: BGI-Shenzhen, Beishan Industrial Zone, Shenzhen 518083, China. Yuanqiang Zou., Address: BGI-Shenzhen, Beishan Industrial Zone, Shenzhen 518083, China. **Contents category:** New taxa - *Firmicutes* and related organisms.

## Abstract

A Gram-staining-negative, non-spore-forming, short, straight rod, non-motile, obligate anaerobic bacterial strain, AF73-05CM02^T^, was isolated from faecal sample of thirty-years old healthy male. Colonies were approximately 0.2mm in diameter, beige and circular after 4 days of incubation on PYG agar under anaerobic conditions at 37°C. Strain AF73-05CM02^T^ grew in a temperature range between 30-42°C and pH range from 6.0 to 8.5, with optimum growth at 37-42°C and pH 7.0. Base on the 16S rRNA gene sequence analysis, strain AF73-05CM02^T^ belongs to the genus *Christensenella* and showed the highest level of sequence similarity with *Christensenella minuta* YIT 12065^T^. The predominant fatty acids of strain AF73-05CM02^T^ were C_10:0_ (7.5%), iso-C_11:0_ (5.6%), C_12:0_ (7.2%), C_14:0_ (46.6%), iso-C_15:0_ (7.4%), C_16:0_ (9.7%) and C_18:1_ ω 9 *c* (6.9%). Acetic acid, formic acid, butyric acid and lactic acid were end products of glucose fermentation. The isolate was negative for catalase, indole production and hydrolysis of gelatin. Genomic relatedness analyses based on average nucleotide identity indicated strain AF73-05CM02^T^ significantly differed from other species of the genus *Christensenella*, showing ANI values under 82.89% with the phylogenetically closest species. The G+C content of the genomic DNA was 52.07 mol% from genome sequence, which differs from that of *Christensenella minuta*. Several physiological biochemical and genotypic properties differentiated the novel bacterial strain from the related species, indicating that the isolate represent a new species of the genus *Christensenella* for which the name *Christensenella intestinihominis* sp. nov. is proposed, with strain AF73-05CM02^T^ (=CGMCC 1.5207^T^ =DSM 103477^T^) being the type strain. The follow study approached the function of cholesterol-lowering for strains AF73-05CM02^T^ and DSM 22067^T^ revealed the two strains exhibit capacity for removing cholesterol with efficiency of 36.6% and 54.3%. Exopolysaccharide production of two strains were 234 and 271 mg/L, respectively.

## Introduction

The human gut is colonised by a large and complex community of microorganisms ranging from 10^13^ to 10^14^ microbial cells (Ventura, 2009; Ghosh, 2013), which is equivalent to 10 times the number of human cells (Bäckhed F, 2005; Jeffery et al., 2016). The microbiome begin to resident the intestinal tract shortly after birth and develops over the first few years (Palmer et al., 2007). The composition of the microbiota is affected by many factors, including the genetic background of the host (Khachatryan et al., 2008; Benson et al., 2010; Fan et al., 2020), the host immune status (Hooper et al., 2012), living condition and daily diet (Turnbaugh et al., 2009; Fujimura et al., 2010). Approximately 90% of gut microbiota were affiliated with the two bacterial phyla, including *Firmicutes* and *Bacteroidetes* (Turnbaugh et al., 2006; Turnbaugh et al., 2008; Tremaroli and Backhed, 2012). *Christensenella minuta* YIT 12065^T^, the type species of genus *Christensenella* within the family *Christensenellaceae*, isolated from human faeces, was first proposed by Morotomi, *et al* (Morotomi et al., 2012). Phylogenetically the isolate formed a novel family-level lineage within the order *Clostridiales* with 86.9–86.1% 16S rRNA gene sequence similarity with the closest relatives. *C. minuta* YIT 12065^T^ was identified as Gram-negative, non-motile, non-spore-forming, short, straight rod with tapered ends and grows anaerobically. The major fatty acids are iso-C_15:0_, C_14:0_ and C_16:0_. LL-diaminopimelic acid is present in the cell wall. The draft genome of *C. minuta* YIT 12065^T^ has been reported previously (Rosa et al., 2017; Coil et al., 2020).The recent research found that strain *C. minuta* as a beneficial bacteria has a significantly effect in protecting against obesity (Goodrich et al., 2014), which might be a novel approach in the treatment of obesity.

Cholesterol is an important basic substance for human body. However, a elevated level of blood cholesterol increases the risk of cardiovascular diseases (CVDs) (Tok and Aslim, 2010; Tsai et al., 2014), which remain the leading cause of deaths worldwide (Ishimwe et al., 2015). In recent years, probiotics have been developed as a non-drug therapy to reduce blood lipids and cholesterol levels and the risk of CVDs (Pan et al., 2010; Tsai et al., 2014). The mechanisms of cholesterol-lowering was proposed for several hypotheses, including deconjugation of bile by bile salt hydrolase activity (Lye et al., 2010a), assimilation and conversion of cholesterol by probiotics (Gilliland et al., 1985; Lye et al., 2010b), and modulating the cholesterol absorption in the intestines of the host (Huang and Zheng, 2010; Yoon et al., 2013).

In the present study, we focus on the polyphasic taxonomic approach for a novel strain, *C. intestinihominis* sp. nov. AF73-05CM02^T^, along with the whole genome sequencing and annotation data, and ivestitgating its cholesterol-lowering property.

## Materials and Method

### Strain isolation

During study the composition of human gut microbiota and construct its taxonomic position by using a polyphasic approach based on phenotypic characteristics and genotypic properties, we isolated a novel *Christensenella*-like stain, designated AF73-05CM02^T^. The fresh faecal sample was collected from a healthy adult residing in Shenzhen, China, and brought back to the lab use for bacteria isolation, The specific location of the studies (GPS coordinates) was 37°35′37″N 114°15′32″E. For cultivation, approximately 1 g fresh faecal was transferred into anaerobic box (Bactron Anaerobic Chamber, Bactron◻-2, shellab, USA) with a gas phase of N_2_/H_2_/CO_2_ (90 : 5 : 5, v/v) and dispersed in 0.1 M PBS (pH 7.0). This suspension containing bacteria was mixed thoroughly and serially diluted and spread onto peptone-yeast extract-glucose (PYG) plates as described previously (Zou et al., 2019). The plate was incubated at 37 °C for 1 week under anaerobic condition. Single colonies were picked and purified by inoculation and subculturing on the same medium. In this study, one of these strains, designated AF73-05CM02^T^ was maintained as a glycerol suspension (20%, w/v) at −80°C. The type strain of genus *Christensenella*, *C. minuta* DSM 22607^T^, procured from the Deutsche Sammlung von Mikroorganismen und Zellkulturen (DSMZ), Braunschweig, Germany, was used as reference strain for phenotypic characterization, genomic comparison and analyses of cell fatty acids and maintained under the same conditions.

### 16S rRNA gene sequencing and phylogenetic analysis

The genomic DNA of strain AF73-05CM02^T^ was prepared from cells harvested from PYG broth using the phenol:chloroform method (Cheng and Jiang, 2006). The 16S rRNA gene was amplified using the universal bacterial primers 27F-1492R (5′-AGAGTTTGATCATGGCTCAG-3′ and 5′-TAGGGTTACCTTGTTACGACTT-3′) and purified as described by Zou *et al.* (Zou et al., 2013). Sequencing was performed by BGI-Shenzhen (Shenzhen, China). The resulting sequence was compared with sequences of type strains retrieved from the EzBioCloud server (Yoon et al., 2017) (https://www.ezbiocloud.net/) using BLAST. Phylogenetic analysis was performed using software package MEGA 7.0 (Tamura et al., 2011) after multiple alignment of sequences data by using CLUSTAL W program (Thompson et al., 1994). Evolutionary phylogenetic trees were constructed using the neighbour-joining method (Saitou and Nei, 1987), maximum-likelihood (Felsenstein, 1981) method and minimum-evolution method (Rzhetsky and Nei, 1993) and bootstrap values were calculated based on 1000 replications.

### Genome sequencing, GC content and genome comparison

For genome comparison of the novel isolate and the closely related species, we conducted genome sequencing and assembly of strain AF73-05CM02^T^. DNA extraction and purity were described above. The draft genome sequence was carried out using a paired-end sequencing strategy with Ion Proton Technology (Life Technologies) at BGI-Shenzhen (Shenzhen, China). The paired-end library had an mean insert size of 500 bp. Reads were assembled using the SOAPdenovo 2 package (Luo et al., 2012). The genomic DNA base content (mol% G+C) was directly calculated from the draft genome data. To determine the DNA relatedness between the isolate and most closely related species, *C. minuta* DSM 22607^T^ and *C. hongkongensis* HKU16^T^ (Lau et al., 2007; Lau et al., 2015), we calculated the average nucleotide identity values (Damodharan et al.), which was thought to be able to corresponds to DNA–DNA hybridization (Goris et al., 2007; Tindall et al., 2010), as described by Kim *et al*. (Kim et al., 2014), following the BLAST-based ANI calculation using the EzGenome web service. ANI values of 95–96% corresponding to 70% DDH has been proposed as a threshold value for species delineation in bacterial taxonomy. The obtained draft genome sequences were annotated using the Rapid Annotation Subsystem Technology (RAST) server (Kanehisa et al., 2016) and KEGG (Aziz et al., 2008) and COG databases (Galperin et al., 2015). A visual genomic comparison across strain AF73-05CM02^T^ and most closely related species was generated with CGView server (Grant and Stothard, 2008) (http://stothard.afns.ualberta.ca/cgview_server/index.html).

### Morphological and growth characteristics

Morphological and cultural characteristics were investigated with strain AF73-05CM02^T^ incubated in PYG medium at 37°C. Morphological observations were examined using both phase contrast microscopy (Olympus BX51, Japan) and transmission electron microscopy (TEM, HITACHI-8100). The Gram reaction, spore formation and presence of flagella were performed by staining using Gram stain kit (Solarbio), spore stain kit (Solarbio) and flagella stain kit (Solarbio) according to the manufacturer’s instructions. Cell motility was examined using semisolid PYG (0.4% agar) (Tittsler RP, 1936). Colony morphology was observed for cultures grown on PYG agar for 4 days at 37°C. Growth at 4, 10, 20, 25, 30, 35, 37, 45 and 50°C was tested on PYG medium to determine the optimal temperature and temperature range for growth. The pH range for growth was evaluated at pH 3.0–10.0 (at interval of 0.5 pH units) by adjusting the pH using the appropriate buffers as described by Sorokin (Sorokin, 2005). Tolerance to NaCl was determined in PYG broth containing different concentrations of NaCl (0-6%, in increments of 1.0%). Bilt tolerance was also measured at different bile salt concentrations (0-5%, in increments of 1.0%) in the PYG broth contained all of the ingredients. All the growth tests of incubation under anraerobic condition for 2 weeks was determined by measuring the OD_600_.

### Physiological and biochemical characteristic

For physiological and biochemical analyse, enzyme activities, hydrolytic activities, utilization of various substrates as sole carbon sources and acid production from different carbohydrates, were carried out for strain AF73-05CM02^T^ comparison with the closely related species, *C. minuta* DSM 22607^T^, using API ZYM, API 20A and API 50CHL systems (bioMérieux, Marcy l’Etoile, France). Sample preparation and test were performed following the directions of the manufacturer’s instructions with incubation at 37°C in an anaerobic condition. For API 50CHL test, CHL broth was supplied with 0.05% cysteine hydrochloride for cells suspension and incubation. Catalase activity was assessed by 3% H_2_O_2_ solution using cells collected from colonies incubated on PYG agar at 37°C for 5 days (Smibert RM, 1994). The isolate and reference type strain were tested under same laboratory conditions.

### Chemotaxonomical characteristic

Chemotaxonomic characteristics of strain AF73-05CM02^T^ and the reference strain were performed by detection of cellular fatty acids and cell wall composition. Strains were cultured on PYG plates at 37°C for 5 days under anaerobic conditions and the fatty acid methyl esters (FAMEs) profile was prepared from lyophilized cells grown in PYG medium by extraction and methylation as described previously (Chen and Dong, 2004). Determination of the fatty acid was analysed by an Agilent HP6890 gas chromatograph and identified using MIDI microbial identification system (M, 1990) and carried out by CGMCC (China General Microbiological Culture Collection Center, Beijing, China). The cell-wall peptidoglycan of strain AF73-05CM02^T^ was performed using wet cell biomass (incubated at 37°C for 5 days on PYG plates) and the amino acid contents in peptidoglycan were determined by TLC as described by Zou, *et al*. (Zou et al., 2013).

### Susceptibility tests and Hemolytic activity

Susceptibility to antibiotics of strain AF73-05CM02^T^ was analysed by the disc diffusion method according to Nizami *et al* (Duran et al., 2012). Antibiotic discs (HANG WEI™, China) were placed on PYG agar plates inoculated with prepared suspensions of the test organisms. The diameter of each zone was measured in millimeters after incubated at 37°C for 5 days. The following antibiotic discs were tested: penicillin (10 ug), ampicillin (10 ug), carbenicillin (100 ug), vancomycin (30), oxacillin (1 ug), piperacillin (100 ug), polymyxin B (300IU), compound sulfamethoxazole (25), furazolidone (300), chloroamphenicol (30) and clindamycin (2). Hemolytic activity was determined in sheep blood agar plates (Guangdong Huankai Microbial Sci&Tech.Co., Ltd.). The plates were incubated under anaerobic conditions for 5 days at 37°C and checked for hemolysis (Pineiro and Stanton, 2007).

### Metabolic end products analysis

The metabolic end products of glucose fermentation, including short-chain fatty acids (SCFAs) and organic acids, were performed using gas chromatograph (GC-7890B, Agilent) equipped with capillary columns and detected using a flame-ionization detector (FID). The capillary column was packed with Agilent 19091N-133HP-INNOWax porapak HP-INNOWax (30m × 0.25mm × 0.25um) for SCFAs detection and Agilent 122-5532G DB-5ms (40m × 0.25mm × 0.25um) for organic acids. The metabolic end products of strain AF73-05CM02^T^ were compared with the closely related species of the genus *Christensenella.*

### The property of Exopolysaccharide (EPS) production

The functional properties of strains AF73-05CM02^T^ and *C. minuta* DSM 22607^T^ were determined by investigating the production of exopolysaccharide (EPS). The EPS were isolated from fermentation solution of two strains using the method as described previously (Mercan et al., 2015). Strains were inoculated in PYG broth at 37°C for 3 days and the cultures were boiled at 100°C for 15 min. The bacterial supernatant was collected after centrifugation at 10,000g for 30 min at 4°C and treated with 80% trichloroacetic acid solution and stirring overnight for precipitating protein. Sample was centrifuged at 10,000g for 30 min at 4°C. The pH of the supernatant was adjusted to 7.0 with 2 M NaOH. The supernatant was then precipitated by adding double-volume chilled ethanol overnight and resuspended in distilled water with gentle heating. EPS was dialyze by 3000 Da dialysis membrane for 24 h at 4°C and washed twice by distilled water. The total EPS production levels was using phenol–sulfuric acid method with glucose as standard (50-500 mg/L) (Dubois, 1956).

### Determination of cholesterol-lowering activity

The capability of strain AF73-05CM02^T^ and the closely related reference strain *C. minuta* DSM 22607^T^ to lower cholesterol was determined according to a modified method of Damodharan *et al* (Damodharan et al., 2015). PYG-CHO broth was prepared with addition of 0.1% (w/v) bile, 0.2% (w/v) sodium thioglycollate and cholesterol dissolved in ethanol at a final concentration of approximately 100μg/ml, w/v. The bacterial culture inoculated with log phase was incubated anaerobically in PYG-CHO at 37°C for 4 days. After incubation, cells were harvested by centrifugation at 10000 × g at 4°C for 10min. The concentration of cholesterol in the supernatant was measured using the o-phthalaldehyde method as described by Rudel and Morris (Rudel, 1973).

Cholesterol-lowering activity from PYG-CHO of each strain broth was calculated in terms of percent cholesterol-lowering as follows:

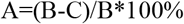

A=% of cholesterol-lowering, B= the concentration of cholesterol in the PYG-CHO, C= the concentration of cholesterol in the supernatant after inoculated with bacteria for 4 days.

## Results and Discussion

### Phylogeny based on 16S rRNA gene sequences

The nearly complete 16S rRNA gene sequence of strain AF73-05CM02^T^ (1,366 bp) was obtained. The closest relatives of the isolate were *C. minuta* DSM 22607^T^, *Catabacter hongkongensis* HKU16^T^ (Lau et al., 2007), *Christensenella massiliensis* Marseille-P2438^T^ (Ndongo et al., 2016b) and *Christensenella timonensis* Marseille-P2437^T^ (Ndongo et al., 2016a) with similarity value of 98.68%, 97.22%, 96.93% and 96.78%, respectively (**Table 1**). Phylogenetic analysis based on the neighbour-joining, maximum-likelihood and minimum-evolution algorithm confirmed that strain AF73-05CM02^T^ is most closely related to *C. minuta* DSM 22607^T^ and formed a tight phylogenetic cluster with 99% bootstrap support (**Figure 1**, **Supplementary Figure S1** and **S2**).

**Table 1.** Levels of 16S rRNA gene sequence similarity and ANI values (in percentages) based on BLAST for strain AF73-05CM02^T^ and the phylogenetically related species. Taxa:1, AF73-05CM02^T^; 2, *C. minuta* DSM 22607^T^; 3, *C. hongkongensis* HKU16^T^; 4, *C. massiliensis* Marseille-P2438^T^; 5, *C. timonensis* Marseille-P2437^T^.

| Strain | Accession no. | 1 | 2* | 3* | 4* | 5* |
| --- | --- | --- | --- | --- | --- | --- |
| **16S rRNA gene sequence similarity (%)** |  |  |  |  |  |  |
| AF73-05CM02<sup>T</sup> | KX078376 | 100 |  |  |  |  |
| *C. minuta* DSM 22607<sup>T</sup> | AB490809 | 98.68 | 100 |  |  |  |
| *C. hongkongensis* HKU16<sup>T</sup> | AB671763 | 97.22 | 96.69 | 100 |  |  |
| *C. massiliensis* Marseille-P2438<sup>T</sup> | LT161898 | 96.93 | 97.51 | 95.99 | 100 |  |
| *C. timonensis* Marseille-P2437<sup>T</sup> | LT223568 | 96.78 | 97.38 | 96.79 | 95.40 | 100 |
| **ANI values (%)** |  |  |  |  |  |  |
| AF73-05CM02<sup>T</sup> | MAIQ00000000 | 100 |  |  |  |  |
| *C. minuta* DSM 22607<sup>T</sup> | NZ_CP029256 | 83.31 | 100 |  |  |  |
| *C. hongkongensis* HKU16<sup>T</sup> | LAYJ00000000 | 73.84 | 75.39 | 100 |  |  |
| *C. massiliensis* Marseille-P2438<sup>T</sup> | LT700187 | 78.00 | 78.01 | 73.28 | 100 |  |
| *C. timonensis* Marseille-P2437<sup>T</sup> | FLKP00000000 | 74.06 | 74.56 | 74.53 | 73.59 | 100 |

\* Data from NCBI and EzBioCloud.

**Figure 1.**
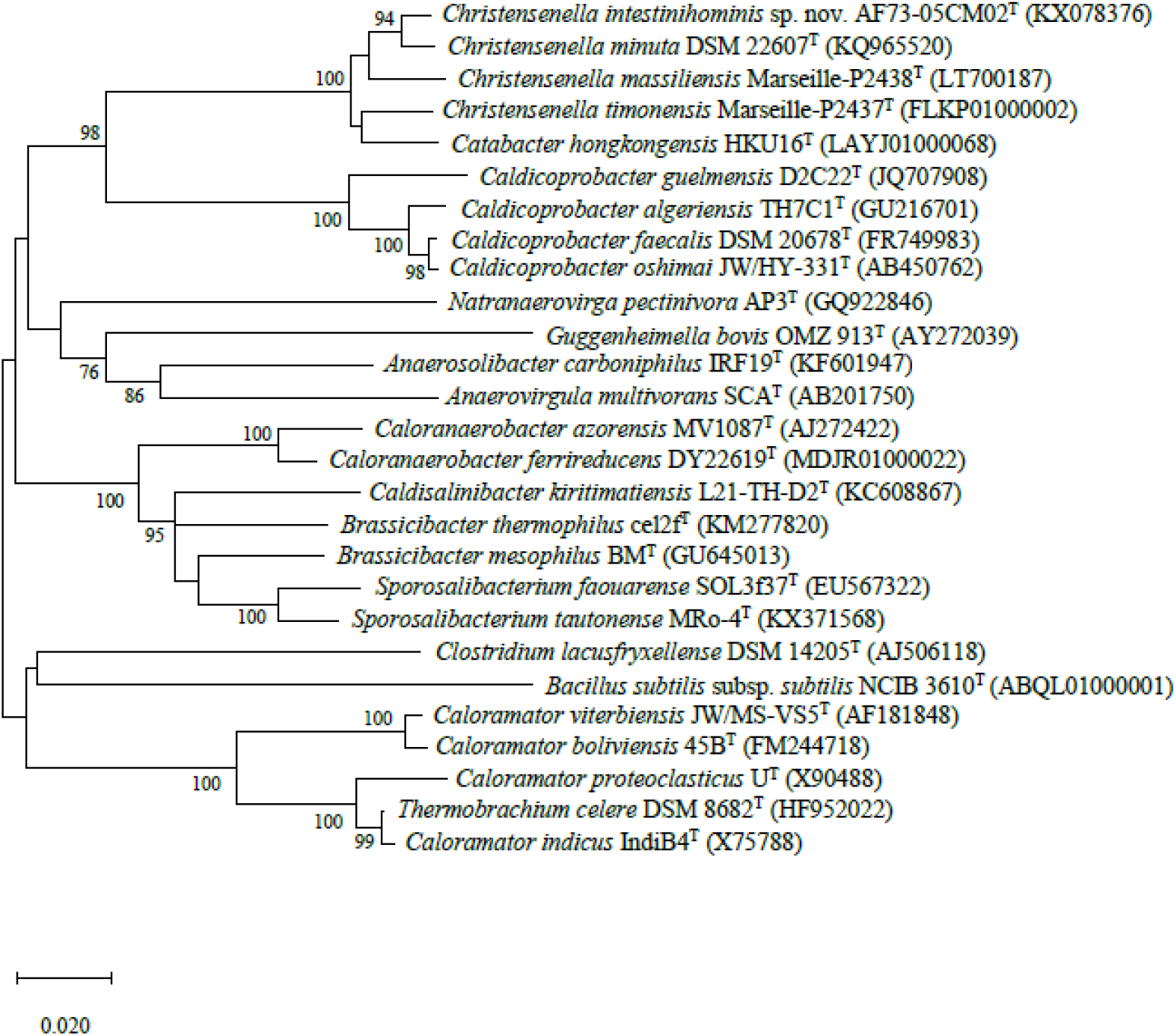
Neighbour-joining phylogenetic tree based on 16S rRNA gene sequences showing the phylogenetic relationships of strains AF73-05CM02^T^ and the representatives of related taxa. *Bacillus subtilis* subsp. *subtilis* NCIB 3610^T^ (ABQL01000001) was used as an out-group. Bootstrap values based on 1000 replications higher than 70% are shown at the branching points. Bar, substitutions per nucleotide position.

### Genome properties

The chromosomes of strain AF73-05CM02^T^ was assembled from 3,145,728 reads resulting a total length of 3,026,655 bp in size and comprised 29 scafolds including 36 contigs. The G+C content of DNA for strain AF73-05CM02^T^ is 52.07 mol% as calculated from the whole-genome sequence. Circular maps of strain AF73-05CM02^T^ in comparison to related species is shown in **Figure 2**. The general features of strain AF73-05CM02^T^ and related species are summarized in **Table 2**.

**Table 2.**
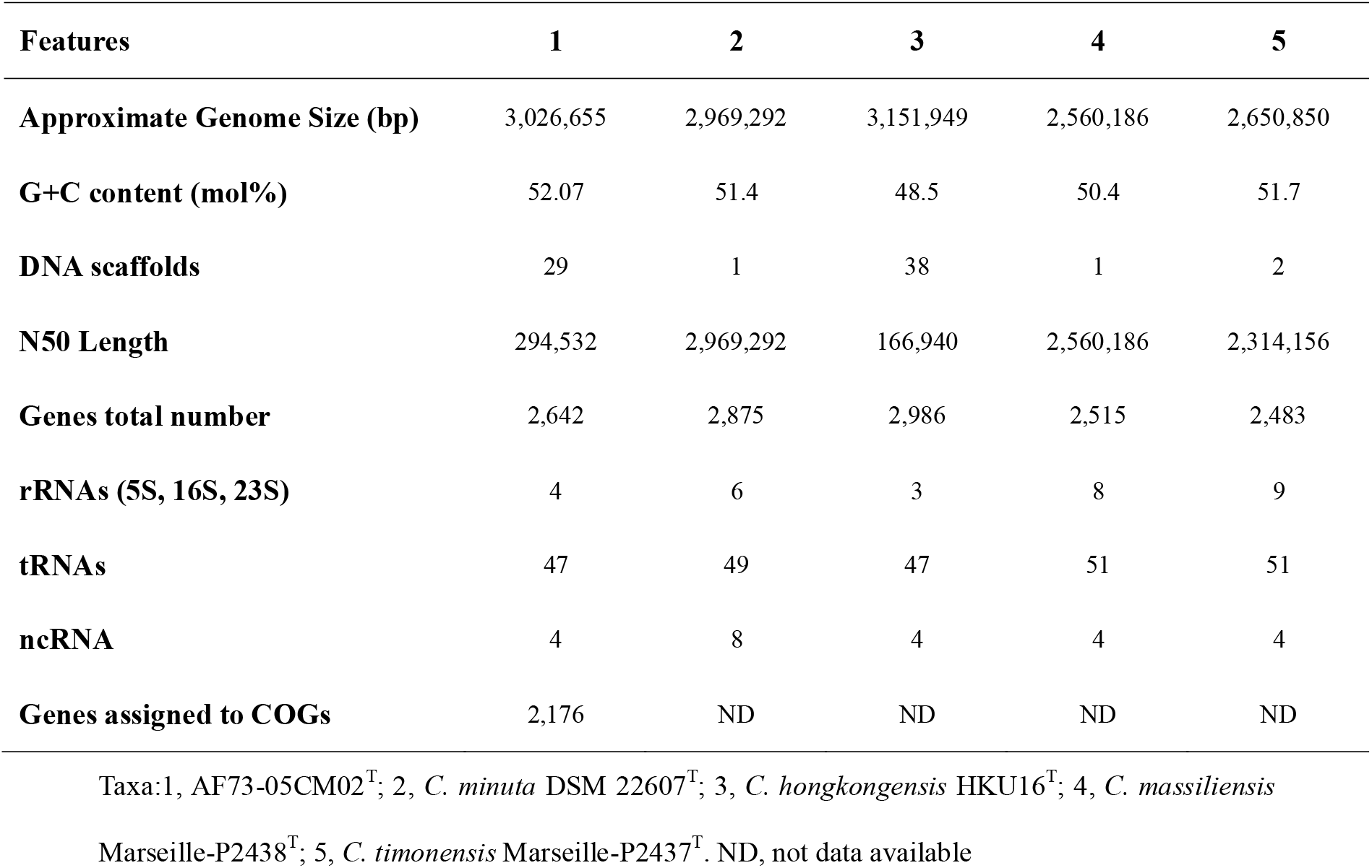
Genome features of *C. intestinihominis* AF73-05CM02^T^ and comparison with closely related species

| Features | 1 | 2 | 3 | 4 | 5 |
| --- | --- | --- | --- | --- | --- |
| **Approximate Genome Size (bp)** | 3,026,655 | 2,969,292 | 3,151,949 | 2,560,186 | 2,650,850 |
| **G+C content (mol%)** | 52.07 | 51.4 | 48.5 | 50.4 | 51.7 |
| **DNA scaffolds** | 29 | 1 | 38 | 1 | 2 |
| **N50 Length** | 294,532 | 2,969,292 | 166,940 | 2,560,186 | 2,314,156 |
| **Genes total number** | 2,642 | 2,875 | 2,986 | 2,515 | 2,483 |
| **rRNAs (5S, 16S, 23S)** | 4 | 6 | 3 | 8 | 9 |
| **tRNAs** | 47 | 49 | 47 | 51 | 51 |
| **ncRNA** | 4 | 8 | 4 | 4 | 4 |
| **Genes assigned to COGs** | 2,176 | ND | ND | ND | ND |

Taxa:1, AF73-05CM02<sup>T</sup>; 2, *C. minuta* DSM 22607<sup>T</sup>; 3, *C. hongkongensis* HKU16<sup>T</sup>; 4, *C. massiliensis* Marseille-P2438<sup>T</sup>; 5, *C. timonensis* Marseille-P2437<sup>T</sup>. ND, not data available

**Figure 2.**
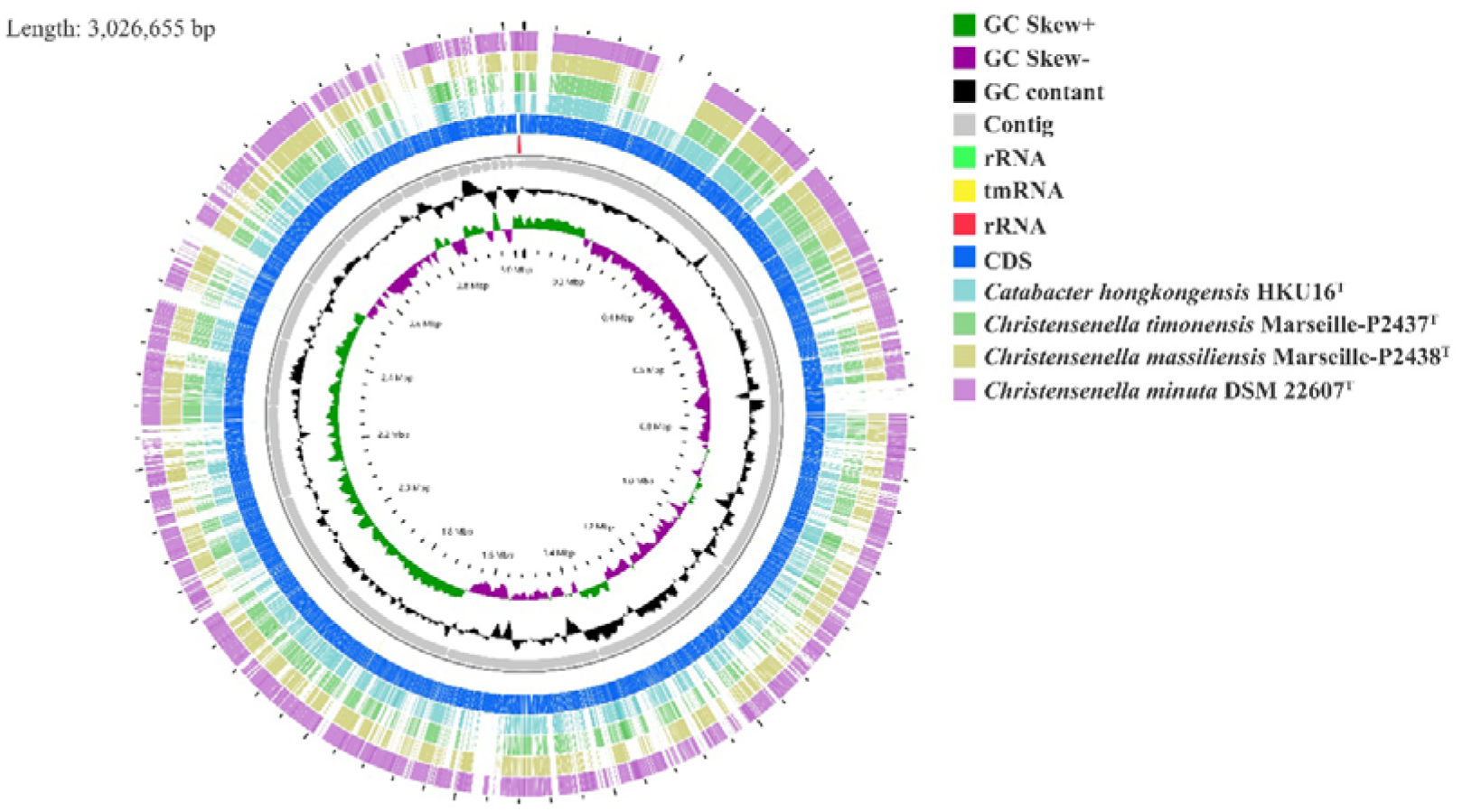
Graphical circular map of the genome from strain *Christensenella intestinihominis* sp. nov. AF73-05CM02^T^, *Christensenella minuta* DSM 22607^T^, *Catabacter hongkongensis* HKU16^T^, *Christensenella massiliensis* Marseille-P2438^T^ and *Christensenella timonensis* Marseille-P2437^T^ using CGView server using default parameters. From inner to outer: Ring 1 and Ring 2, G+C positive skew (green) and G+C negative skew (purple); Ring 3, GC% content; Ring 4-Ring 8, Contig, rRNA, tmRNA, rRNA and CDS from AF73-05CM02^T^; Ring 9, *Catabacter hongkongensis* HKU16^T^; Ring 10, *Christensenella timonensis* Marseille-P2437^T^; Ring 11, *Christensenella massiliensis* Marseille-P2438^T^; Ring 12, *Christensenella minuta* DSM 22607^T^.

Among the 2,642 annotated genes in the *C. intestinihominis* AF73-05CM02^T^ genome, 2,176 genes with specific functions were assigned to COGs. The distribution of genes into COGs functional classification was presented in **Figure 3** and **Supplementary Table S1**, revealed that E (Amino acid transport and metabolism), G (Carbohydrate transport and metabolism), M (Cell wall/membrane/envelope biogenesis), C (Energy production and conversion), R (General function prediction only), T (Signal transduction mechanisms), K (Transcription) and J (Translation, ribosomal structure and biogenesis) were abundant categories. For compared the Individual predicted coding sequences of strain AF73-05CM02^T^ with *C. minuta* DSM 22607^T^ by RAST annotation, we found there were 10-11 RAST-annotated genes associated with diaminopimelic acid synthesis, 31-37 genes associated with metabolism of polar lipids, 14-16 genes associated with metabolism of polyamines, 4-5 genes associated with teichoic and lipoteichoic acids biosynthesis, and 3 genes associated with lipopolysaccharides biosynthesis present in the genomes (**Table 3** and **Supplementary Table S2**). The number and kind of genes associated with diaminopimelic acid, polar lipids, polyamines and teichoic and lipoteichoic acids biosynthesis make strain AF73-05CM02^T^ distinguishable from the reference species, *C. minuta* DSM 22607^T^.

**Figure 3.**
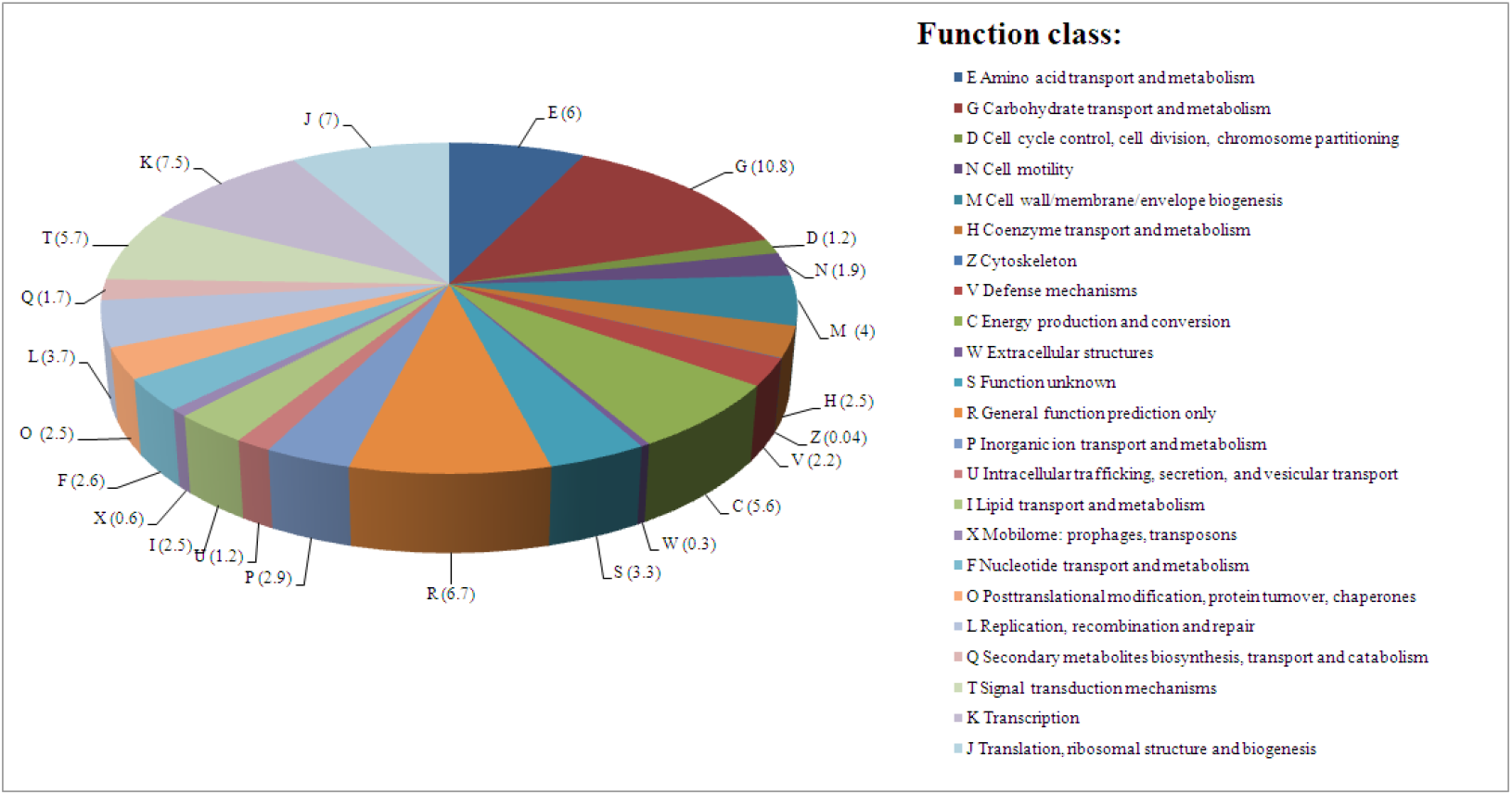
The distribution of the genes associated with the COG functional categories in strain AF73-05CM02^T^. The number of genes is shown in parentheses.

**Table 3.** Number of genes associated with biosynthetic pathway from whole genome sequences of strain AF73-05CM02^T^ and *C. minuta* DSM 22607^T^ identified by RAST. Numbers of genes identified for mycolic acids and quinines (benzoquinones and naphthoquinones) were zero for all taxa studied.

| Genes responsible for biosynthesis | AF73-05CM02<sup>T</sup> | *C. minuta* DSM 22607<sup>T</sup> |
| --- | --- | --- |
| **Diaminopimelic acid** | 10 | 11 |
| **Polar lipids** | 31 | 37 |
| **Polyamines** | 14 | 16 |
| **Teichoic and lipoteichoic acids** | 4 | 5 |
| **Lipopolysaccharides** | 3 | 3 |

In order to further distinguish strain AF73-05CM02^T^ from the phylogenetically related species, the genome comparison was determined using BLAST average nucleotide identities (ANIb). The ANI values between strain AF73-05CM02^T^ and related reference species, *C. minuta* DSM 22607^T^, *C. hongkongensis* HKU16^T^, *C. massiliensis* Marseille-P2438^T^ and *C. timonensis* Marseille-P2437^T^ were calculated as 83.51%, 78.92%, 79.66% and 78.76%, respectively (**Table 1**). The ANI values of strain AF73-05CM02^T^ with the related species were significantly below the cutoff of 95–96%, which is proposed as a threshold value for species delineation in bacterial taxonomy (Goris et al., 2007), indicating that strain AF73-05CM02^T^ is a distinct genomic species and should be classified as a representative of a novel species.

### Phenotypic features

Strain AF73-05CM02^T^ was an obligate anaerobic and Gram-stain-negative bacterium. Cells were approximately 0.5μm in width and 1.0–2.0μm in length and occurring singly or in short chains. Under phase contrast microscope, cells were non-spore-forming, flagella were not observed. The bacteria formed punctiform colonies (approximately 0.2mm in diameter) with circular and beige after 4 days of growth at 37°C on PYG agar under anaerobic conditions. The growth temperature was from 30-42°C, with the optimum around 37-42°C, while no growth was observed below 30°C or at 45°C. Growth occured at pH values from 6.0 to 8.5, with optimum growth between 6.5 and 7.0. The strain tolerated salt concentrations up to 2% (w/v) NaCl and bile up to 0.3%. The cells were catalase-negative. The physiological and biochemical comparison of strain AF73-05CM02^T^ and related strain was carried out using API 20A, API 50CHL and API ZYM tests, the result were summarized in the species description and the differences of selected characteristics with the reference strain are given in **Table 4**. All the results of enzymatic characteristics and carbon source assimilation from API ZYM, API 20A and API 50CHL test are presented in **Supplementary Table S3** and **Supplementary Table S4.**

**Table 4.**
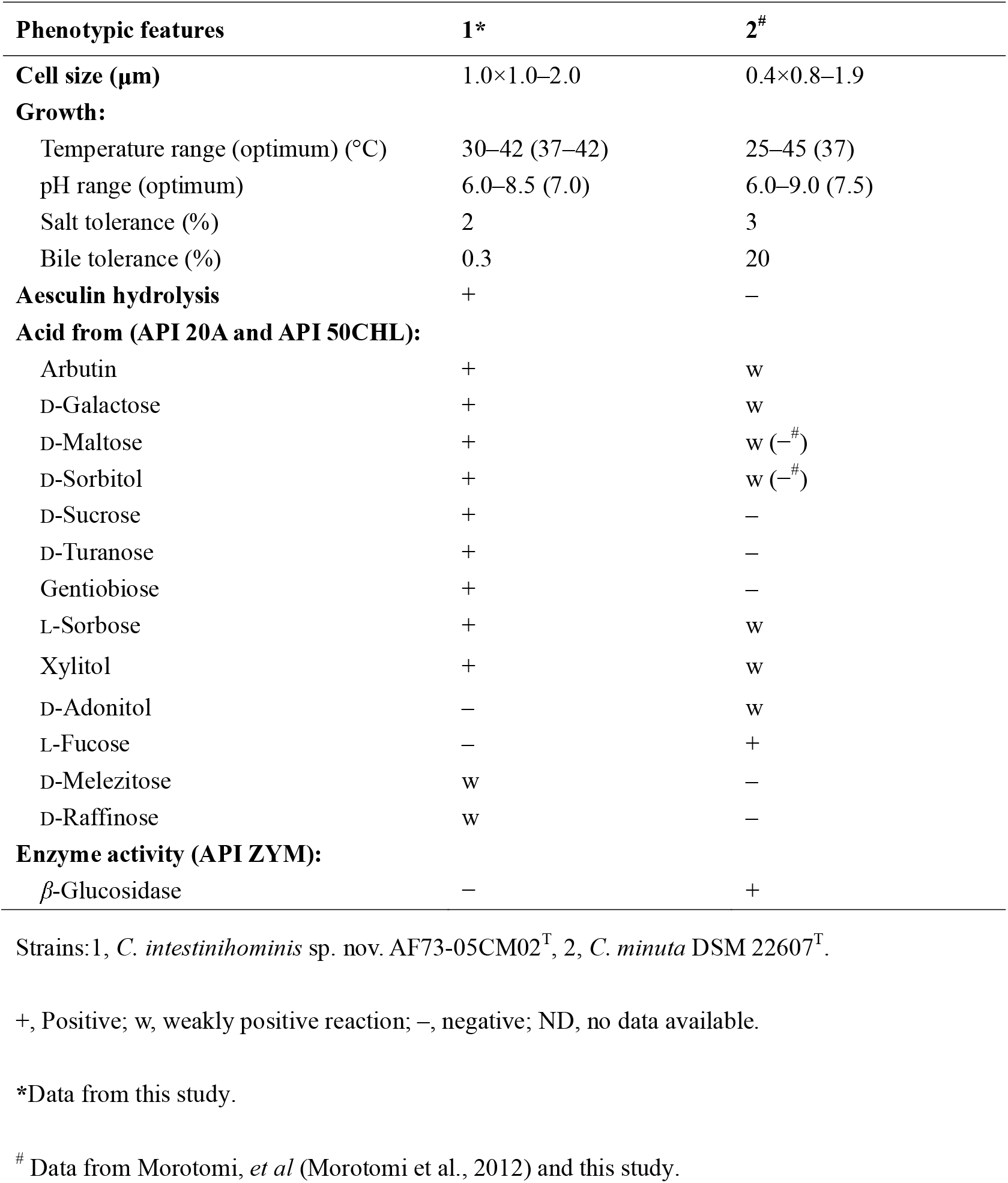
Comparison of phenotypic features between strain *C. intestinihominis* AF73-05CM02^T^ and the closest related reference strain, *C. minuta* DSM 22607^T^.

Chemotaxonomic characteristics of strain AF73-05CM02^T^ were consistent with the results of the reference strain that were performed under identical conditions, confirming that the novel isolate belongs to the genus *Christensenella*. The cellular fatty acid composition of strain AF73-05CM02^T^ and DSM 22607^T^ are presented in **Table 5**, and the dominant fatty acids (representing > 5% of the total) for strain AF73-05CM02^T^ were C_10:0_ (7.5%), iso-C_11:0_ (5.6%), C_12:0_ (7.2%), C_14:0_ (46.6%), iso-C_15:0_ (7.4%), C_16:0_ (9.7%) and C_18:1_ ω9*c* (6.9%). The higher amount of C_14:0_ and less amount of iso-C_15:0_ and C_16:0_ significantly differentiated strain AF73-05CM02^T^ from the reference strains. The cell-wall diamino acid was LL-diaminopimelic acid.

**Table 5.** Cellular fatty acid composition of strain AF73-05CM02^T^ and closely related species, DSM 22607^T^. Strains: 1, AF73-05CM02^T^; 2, *C. minuta* DSM 22607^T^; Data were obtained in this study. Numbers represent percentages of the total fatty acids. Only fatty acids amounting 1% or higher are shown. t, traces (<1%).

| Fatty acids | 1 | 2 |
| --- | --- | --- |
| C<sub>10:0</sub> | 7.5 | 8.6 |
| C<sub>12:0</sub> | 7.2 | 1.1 |
| C<sub>14:0</sub> | 46.6 | 13.0 |
| C<sub>14:0</sub> 2OH | t | 1.3 |
| C<sub>16:0</sub> | 9.7 | 21.1 |
| C<sub>18:1</sub> *ω*9*c* | 6.9 | 6.8 |
| C<sub>18:1</sub> *ω*7*c* | t | 3.9 |
| C<sub>18:0</sub> | 1.8 | 3.7 |
| Iso-C<sub>11:0</sub> | 5.6 | 2.9 |
| Iso-C<sub>15:0</sub> | 7.4 | 27.4 |
| Anteiso-C<sub>11:0</sub> | t | 1.3 |
| Anteiso-C<sub>13:0</sub> | t | 2.4 |
| Anteiso-C<sub>15:0</sub> | 1.3 | 3.2 |
| Iso-C<sub>17:1</sub> I/anteiso B | 4.7 | 1.7 |
| Antei-C<sub>18:0</sub> /C<sub>18:2</sub> *ω*6,9*c* | t | 1.7 |

For Susceptibility tests, strain AF73-05CM02^T^ was resistant to oxacillin and compound sulfamethoxazole, but sensitive to penicillin, ampicillin, carbenicillin, piperacillin, vancomycin, polymyxin B, furazolidone, chloroamphenicol and clindamycin (**Supplementary Table S5**). The hemolytic activity of the cells was not founded.

Metabolic end products from glucose for strain AF73-05CM02^T^ and DSM 22607 ^T^ are shown in **Supplementary Table 6**. Acetic acid, formic acid, butyric acid and lactic acid were the major end products (>1 mmol/L) for strain AF73-05CM02^T^.

We found strain AF73-05CM02^T^ can be clearly differentiated from *C. minuta* DSM 22607^T^ based on a lot of phenotypic and genotypic characteristics and ANI values obtained in this study can separate this isolate from related species, which suggest that the strain AF73-05CM02^T^ represents a novel species of the genus *Christensenella.* Therefore, we propose AF73-05CM02^T^ (=CGMCC 1.5207^T^ =DSM 103477^T^) as the type strain of *Christensenella intestinihominis* sp. nov.

### EPS production

EPS produced by probiotics have several biologically beneficial functions on the host, such as improve the viscosity of the lactic acid bacteria fermented product (Li et al., 2014) and have significant roles on colonization, stress resistance and adhesion (Delcour et al., 1999). Furthermore, it has been suggested EPS may have probiotics properties on immune modulation and antioxidative effects (Welman and Maddox, 2003; Fanning et al., 2012). In the present research, both test strains, *C. intestinihominis* AF73-05CM02^T^ and *C. minuta* DSM 22607^T^, were capable of producing EPS with amount of 234 and 271 mg/L, respectively.

### Removal of Cholesterol

The test the cholesterol-lowering activity was determined in PYG-CHO broth added with bile. Both strain AF73-05CM02^T^ and *C. minuta* DSM 22607^T^ showed a capacity for removing cholesterol from PYG-CHO broth. After incubated in PYG-CHO at 37°C for 4 days, the amount of cholesterol in medium were reduced with efficiency of 36.6% and 54.3% by AF73-05CM02^T^ and *C. minuta* DSM 22607^T^, respectively. The control sample, containing no cultures, demonstrated no cholesterol removal, as expected. The mechanisms of cholesterol-lowering by probiotics from *in vitro* experiments have been reported including many hypotheses, such as deconjugated bile acids via bile salt hydrolase activity, adsorption to cellular surface and conversion by probiotics (Ishimwe et al., 2015). In the previous study, *in vivo* experiment of cholesterol-lowering showed the probiotics have a useful and safe effect on modulating the serum-lipid profile and reducing the host cholesterol level (Pan et al., 2010). A high cholesterol level as a main production of obesity can increase the risk of CVDs. The genus *Christensenella* as a especially common microorganism has been founded in lean people and showed a high abundence population in the gut (Goodrich et al., 2014), suggesting *Christensenella* has a potential function in protecting against obesity. Further studies will be required to focus on the cholesterol reducing properties *in vitro* and *in vivo* and reveal the mechanism of lose weight for genus *Christensenella*.

### Description of *Christensenella intestinihominis* sp. nov

*Christensenella intestinihominis* (in.tes.ti.ni.ho′mi.nis. L. gen. n. *intestini* of the intestine; L. gen. n. *hominis* of a human being; N.L. gen. n. *intestinihominis* of the human intestine).

Cells are Gram-stain-negative, obligately anaerobic, non-motile and short rods (1.0×1.0–2.0μm) isolated from a faecal sample collected from a healthy adult. Colonies on PYG agar are 0.2mm in diameter and punctiform with circular and beige after 4 days of growth at 37°C. Growth occurs at temperatures from 30-42°C, with the optimum around 37-42°C. The pH range is from 6.0 to 8.5, with optimum between 6.5 and 7.0. Able to grow in the presence of up to 2.0% (w/v) NaCl and 0.3% bile (w/v). Major end products of metabolism of glucose are Acetic acid, formic acid, butyric acid and lactic acid. The cells exhibit resistance to oxacillin and compound sulfamethoxazole, but are sensitive to penicillin, ampicillin, carbenicillin, piperacillin, vancomycin, polymyxin B, furazolidone, chloroamphenicol, and clindamycin. The predominant cellular fatty acids are C_10:0_, iso-C_11:0_, C_12:0_, C_14:0_, iso-C_15:0_, C_16:0_ and C_18:1_ ω9*c*. The diagnostic cell-wall diamino acid is LL-diaminopimelic acid.

In API 20A and API 50CHL, the isolate was positive for utilization arbutin, D-arabinose, D-fructose, D-fucose, D-galactose, D-glucose, D-lyxose, D-ribose, D-sorbitol, D-sucrose, D-tagatose, D-turanose, D-xylose, gentiobiose, L-arabinose, L-rhamnose, L-sorbose, methyl-*β*-D-xylopyranoside, salicin and xylitol, weakly reactions for D-maltose, D-mannose, D-melezitose, D-raffinose, erythritol, L-xylose and salicin, and negative for amygdalin, cellobiose, D-adonitol, D-arabitol, D-lactose, D-mannitol, D-melibiose, D-trehalose, dulcitol, gluconate, glycerol, glycogen, inositol, inulin, L-arabitol, L-fucose, methyl-D-glucopyranoside, methyl-*α*-D-mannopyranoside, *N*-acetyl-glucosamine, 2-ketogluconate and 5-ketogluconate. Indole is not formed. Esculin can be degraded, but gelatin is not hydrolysed. Catalase is negative.

Results obtained from API ZYM showed positive enzymatic activity for naphthol-AS-BI-phosphohydrolase and negative for alkaline phosphatase, esterase (C4), esterase lipase (C8), lipase (C14), leucine arylamidase, valine arylamidase, cystine arylamidase, trypsin, *α*-chymotrypsin, acid phosphatase, *α*-galactosidase, *β*-galactosidase, *β*-glucuronidase, *α*-glucosidase, *β*-glucosidase, *N*-acetyl-*β*-glucosaminidase, *α*-mannosidase and *β*-fucosidase.

In the result of RAST annotation, 11 genes/proteins are accociated with biosynthesis of DAP, including 4-hydroxy-tetrahydrodipicolinate reductase (EC 1.17.1.8), 4-hydroxy-tetrahydrodipicolinate synthase (EC 4.3.3.7), aspartate-semialdehyde dehydrogenase (EC 1.2.1.11), aspartokinase (EC 2.7.2.4), diaminopimelate decarboxylase (EC 4.1.1.20), diaminopimelate epimerase (EC 5.1.1.7), L, L-diaminopimelate aminotransferase (EC 2.6.1.83), *N*-acetyl-L, L-diaminopimelate deacetylase (EC 3.5.1.47), *N*-succinyl-L, L-diaminopimelate desuccinylase (EC 3.5.1.18), UDP-*N*-acetylmuramoylalanyl-D-glutamate-2, 6-diaminopimelate ligase (EC 6.3.2.13) and UDP-*N*-acetylmuramoylalanyl-D-glutamyl-2, 6-diaminopimelate-D-alanyl-D-alanine ligase (EC 6.3.2.10). 31 genes/proteins are accociated with biosynthesis of polar lipids, including 1-acyl-sn-glycerol-3-phosphate acyltransferase (EC 2.3.1.51), acyl carrier protein (4 copies), acyl-phosphate:glycerol-3-phosphate O-acyltransferase PlsY, alcohol dehydrogenase (EC 1.1.1.1) (8 copies), acetaldehyde dehydrogenase (EC 1.2.1.10) (2 copies), aldehyde dehydrogenase (EC 1.2.1.3), aldehyde dehydrogenase B (EC 1.2.1.22), cardiolipin synthetase (EC 2.7.8.-), CDP-diacylglycerol-glycerol-3-phosphate 3-phosphatidyltransferase (EC 2.7.8.5), diacylglycerol kinase (EC 2.7.1.107), dihydroxyacetone kinase family protein, glycerate kinase (EC 2.7.1.31), glycerol kinase (EC 2.7.1.30) (2 copies), glycerol-1-phosphate dehydrogenase [NAD(P)] (EC 1.1.1.261) (2 copies), glycerol-3-phosphate dehydrogenase (EC 1.1.5.3), glycerol-3-phosphate dehydrogenase [NAD(P)^+^] (EC 1.1.1.94), phosphate:acyl-ACP acyltransferase PlsX and phosphatidate cytidylyltransferase (EC 2.7.7.41). 14 genes/proteins are accociated with biosynthesis of polyamines, including agmatine deiminase (EC 3.5.3.12), agmatine/putrescine antiporter, associated with agmatine catabolism (2 copies), arginine decarboxylase (EC 4.1.1.19) / Lysine decarboxylase (EC 4.1.1.18), carbamate kinase (EC 2.7.2.2), carboxynorspermidine dehydrogenase, putative (EC 1.1.1.-), putrescine carbamoyltransferase (EC 2.1.3.6), putrescine transport ATP-binding protein PotA (TC 3.A.1.11.1), S-adenosylmethionine decarboxylase proenzyme (EC 4.1.1.50), prokaryotic class 1A and spermidine putrescine ABC transporter permease component PotB (TC 3.A.1.11.1), spermidine putrescine ABC transporter permease component potC (TC_3.A.1.11.1) (2 copies), spermidine synthase (EC 2.5.1.16) and transcriptional regulator, MerR family, near polyamine transporter. 4 genes/proteins are accociated with biosynthesis of teichoic and lipoteichoic acids, including 2-C-methyl-D-erythritol 4-phosphate cytidylyltransferase (EC 2.7.7.60), teichoic acid export ATP-binding protein TagH (EC 3.6.3.40), teichoic acid translocation permease protein TagG and undecaprenyl-phosphate *N*-acetylglucosaminyl 1-phosphate transferase (EC 2.7.8.-). 3 genes/proteins are accociated with biosynthesis of lipopolysaccharides, including lipopolysaccharide biosynthesis protein RffA (2 copies), lipopolysaccharide cholinephosphotransferase LicD1 (EC 2.7.8.-) and HtrA protease/chaperone protein. There are no genes responsible for biosynthesis of respiratory lipoquinones or mycolic acids.

The type strain AF73-05CM02^T^ (=CGMCC 1.5207^T^ =DSM 103477^T^) was isolated from the faecal samples of a healthy adult residing in Shenzhen, China (37°35′37″N, 114°15′32″E). The DNA G+C content of strain AF73-05CM02^T^ is 52.07 mol% calculated from the genome sequence. The genome size is 3.02Mbp.

## Data Availability Statement

The GenBank/EMBL/DDBJ accession number for the 16S rRNA gene sequence of *Christensenella intestinihominis* AF73-05CM02^T^ are KX078376. The draft genome of *C. intestinihominis* AF73-05CM02^T^ have been deposited at DDBJ/EMBL/GenBank under the accession numbers MAIQ00000000. The data that support the findings of this study have also been deposited into CNGB Sequence Archive (CNSA) (Guo et al., 2020) of China National GeneBank DataBase (CNGBdb) (Chen et al., 2020) with accession number CNPhis0003415.

## Author Contributions

Conceived and designed the experiments:Y.Z. and L.X. Performed the experiments: Y.Z., W.X., M.L. and Y.D. Analyzed the data: Y.Z., L.X., G.L., and X.L. Contributed reagents/materials/analysis tools: Y.Z., W.X., M.L. and Y.D. Wrote the paper: YZ.

## Acknowledgements

We thank the colleagues at BGI-Shenzhen for sample collection, and discussions, and China National Genebank (CNGB) Shenzhen for DNA extraction, library construction, sequencing.

## Supplementary Material

**Supplementary Table S1. Number of genes associated with general COG functional categories in the genome of *C. intestinihominis* AF73-05CM02^T^ and *C. minuta* DSM 22607^T^.**

**Supplementary Table S2. The specific genes/protein related to biosynthesis of DAP, polar lipids, polyamines and lipoteichoic and teichoic acids and their positions in the genome in comparasion of strain AF73-05CM02^T^ and *C. minuta* DSM 22607^T^ identified by Rapid Annotation Subsystem Technology (RAST).**

**Supplementary Table S3. Enzymatic characteristics of strain AF73-05CM02^T^ from API ZYM test.**

**Supplementary Table S4. Carbon source assimilation of strain AF73-05CM02^T^ from API 20A and API 50CHL test.**

**Supplementary Table S5. Antibiotic sensitivity of strain AF73-05CM02^T^.**

**Supplementary Table S6. Metabolic end products from glucose for strain AF73-05CM02^T^ and *C. minuta* DSM 22607^T^.**

**Supplementary Figure S1. Maximum-likelihood phylogenetic tree based on 16S rRNA gene sequences showing the phylogenetic relationships of strains AF73-05CM02^T^ and the representatives of related taxa.** *Bacillus subtilis subsp. subtilis* NCIB 3610^T^ (ABQL01000001) was used as an out-group. Bootstrap values based on 1000 replications higher than 70% are shown at the branching points. Bar, substitutions per nucleotide position.

**Supplementary Figure S2. Minimum-evolution phylogenetic tree based on 16S rRNA gene sequences showing the phylogenetic relationships of strains AF73-05CM02^T^ and the representatives of related taxa.** *Bacillus subtilis subsp. subtilis* NCIB 3610^T^ (ABQL01000001) was used as an out-group. Bootstrap values based on 1000 replications higher than 70% are shown at the branching points. Bar, substitutions per nucleotide position.

**Supplementary Figure S3. Certification. Deposit certification of CGMCC.**

**Supplementary Figure S4. Certification. Deposit certification of DSMZ.**

## Reference

Aziz, R.K., Bartels, D., Best, A.A., DeJongh, M., Disz, T., Edwards, R.A., et al. (2008). The RAST Server: rapid annotations using subsystems technology. BMC Genomics 9, 75. doi: 10.1186/1471-2164-9-75.

Bäckhed F, L.R., Sonnenburg JL, Peterson DA, Gordon JI. (2005). Host-Bacterial Mutualism in the Human Intestine. Science 307(5717), 1915–1920. doi: 10.1126/science.1104816.

Benson, A.K., Kelly, S.A., Legge, R., Ma, F., Low, S.J., Kim, J., et al. (2010). Individuality in gut microbiota composition is a complex polygenic trait shaped by multiple environmental and host genetic factors. Proc Natl Acad Sci U S A 107(44), 18933–18938. doi: 10.1073/pnas.1007028107.

Chen, F.Z., You, L.J., Yang, F., Wang, L.N., Guo, X.Q., Gao, F., et al. (2020). CNGBdb: China National GeneBank DataBase. Yi Chuan 42(8), 799–809. doi: 10.16288/j.yczz.20-080.

Chen, S., and Dong, X. (2004). *Acetanaerobacterium elongatum* gen. nov., sp. nov., from paper mill waste water. Int J Syst Evol Microbiol 54(Pt 6), 2257–2262. doi: 10.1099/ijs.0.63212-0.

Cheng, H.R., and Jiang, N. (2006). Extremely rapid extraction of DNA from bacteria and yeasts. Biotechnol Lett 28(1), 55–59. doi: 10.1007/s10529-005-4688-z.

Coil, D.A., Jospin, G., and Eisen, J.A. (2020). Draft Genome Analysis of *Christensenella minuta* DSM 22607, exhibiting an unusual expansion of transporter homologs of unknown function. J Genomics 8, 25–29. doi: 10.7150/jgen.43162.

Damodharan, K., Lee, Y.S., Palaniyandi, S.A., Yang, S.H., and Suh, J.W. (2015). Preliminary probiotic and technological characterization of *Pediococcus pentosaceus* strain KID7 and in vivo assessment of its cholesterol-lowering activity. Front Microbiol 6, 768. doi: 10.3389/fmicb.2015.00768.

Delcour, J., Ferain, T., Deghorain, M., Palumbo, E., and Hols, P. (1999). The biosynthesis and functionality of the cell-wall of lactic acid bacteria. Antonie Van Leeuwenhoek 76(1-4), 159–184.

Dubois, M., Gilles, K. A., Hamilton, J. K., Rebers, P. A., Smith, F. (1956). Colorimetric method for determination of sugars and related substances. Analytical Chemistry 28, 350–356.

Duran, N., Ozer, B., Duran, G.G., Onlen, Y., and Demir, C. (2012). Antibiotic resistance genes & susceptibility patterns in staphylococci. Indian J Med Res 135, 389–396.

Fan, P., Bian, B., Teng, L., Nelson, C.D., Driver, J., Elzo, M.A., et al. (2020). Host genetic effects upon the early gut microbiota in a bovine model with graduated spectrum of genetic variation. ISME J 14(1), 302–317. doi: 10.1038/s41396-019-0529-2.

Fanning, S., Hall, L.J., Cronin, M., Zomer, A., MacSharry, J., Goulding, D., et al. (2012). Bifidobacterial surface-exopolysaccharide facilitates commensal-host interaction through immune modulation and pathogen protection. Proc Natl Acad Sci U S A 109(6), 2108–2113. doi: 10.1073/pnas.1115621109.

Felsenstein, J. (1981). Evolutionary trees from DNA sequences: a maximum likelihood approach. J Mol Evol 17(6), 368–376.

Fujimura, K.E., Slusher, N.A., Cabana, M.D., and Lynch, S.V. (2010). Role of the gut microbiota in defining human health. Expert Rev Anti Infect Ther 8(4), 435–454. doi: 10.1586/eri.10.14.

Galperin, M.Y., Makarova, K.S., Wolf, Y.I., and Koonin, E.V. (2015). Expanded microbial genome coverage and improved protein family annotation in the COG database. Nucleic Acids Res 43(Database issue), D261–269. doi: 10.1093/nar/gku1223.

Ghosh, A.R. (2013). Appraisal of microbial evolution to commensalism and pathogenicity in humans. Clin Med Insights Gastroenterol 6, 1–12. doi: 10.4137/CGast.S11858.

Gilliland, S.E., Nelson, C.R., and Maxwell, C. (1985). Assimilation of cholesterol by *Lactobacillus acidophilus*. Appl Environ Microbiol 49(2), 377–381.

Goodrich, J.K., Waters, J.L., Poole, A.C., Sutter, J.L., Koren, O., Blekhman, R., et al. (2014). Human genetics shape the gut microbiome. Cell 159(4), 789–799. doi: 10.1016/j.cell.2014.09.053.

Goris, J., Konstantinidis, K.T., Klappenbach, J.A., Coenye, T., Vandamme, P., and Tiedje, J.M. (2007). DNA-DNA hybridization values and their relationship to whole-genome sequence similarities. Int J Syst Evol Microbiol 57(Pt 1), 81–91. doi: 10.1099/ijs.0.64483-0.

Grant, J.R., and Stothard, P. (2008). The CGView Server: a comparative genomics tool for circular genomes. Nucleic Acids Res 36(Web Server issue), W181–184. doi: 10.1093/nar/gkn179.

Guo, X., Chen, F., Gao, F., Li, L., Liu, K., You, L., et al. (2020). CNSA: a data repository for archiving omics data. Database (Oxford) 2020. doi: 10.1093/database/baaa055.

Hooper, L.V., Littman, D.R., and Macpherson, A.J. (2012). Interactions between the microbiota and the immune system. Science 336(6086), 1268–1273. doi: 10.1126/science.1223490.

Huang, Y., and Zheng, Y. (2010). The probiotic *Lactobacillus acidophilus* reduces cholesterol absorption through the down-regulation of Niemann-Pick C1-like 1 in Caco-2 cells. Br J Nutr 103(4), 473–478. doi: 10.1017/S0007114509991991.

Ishimwe, N., Daliri, E.B., Lee, B.H., Fang, F., and Du, G. (2015). The perspective on cholesterol-lowering mechanisms of probiotics. Mol Nutr Food Res 59(1), 94–105. doi: 10.1002/mnfr.201400548.

Jeffery, I.B., Lynch, D.B., and O’Toole, P.W. (2016). Composition and temporal stability of the gut microbiota in older persons. ISME J 10(1), 170–182. doi: 10.1038/ismej.2015.88.

Kanehisa, M., Sato, Y., Kawashima, M., Furumichi, M., and Tanabe, M. (2016). KEGG as a reference resource for gene and protein annotation. Nucleic Acids Res 44(D1), D457–462. doi: 10.1093/nar/gkv1070.

Khachatryan, Z.A., Ktsoyan, Z.A., Manukyan, G.P., Kelly, D., Ghazaryan, K.A., and Aminov, R.I. (2008). Predominant role of host genetics in controlling the composition of gut microbiota. PLoS One 3(8), e3064. doi: 10.1371/journal.pone.0003064.

Kim, M., Oh, H.S., Park, S.C., and Chun, J. (2014). Towards a taxonomic coherence between average nucleotide identity and 16S rRNA gene sequence similarity for species demarcation of prokaryotes. Int J Syst Evol Microbiol 64(Pt 2), 346–351. doi: 10.1099/ijs.0.059774-0.

Lau, S.K., McNabb, A., Woo, G.K., Hoang, L., Fung, A.M., Chung, L.M., et al. (2007). *Catabacter hongkongensis* gen. nov., sp. nov., isolated from blood cultures of patients from Hong Kong and Canada. J Clin Microbiol 45(2), 395–401. doi: 10.1128/JCM.01831-06.

Lau, S.K., Teng, J.L., Huang, Y., Curreem, S.O., Tsui, S.K., and Woo, P.C. (2015). Draft Genome Sequence of *Catabacter hongkongensis* Type Strain HKU16T, Isolated from a Patient with Bacteremia and Intestinal Obstruction. Genome Announc 3(3). doi: 10.1128/genomeA.00531-15.

Li, W., Ji, J., Chen, X., Jiang, M., Rui, X., and Dong, M. (2014). Structural elucidation and antioxidant activities of exopolysaccharides from *Lactobacillus helveticus* MB2-1. Carbohydr Polym 102, 351–359. doi: 10.1016/j.carbpol.2013.11.053.

Luo, R., Liu, B., Xie, Y., Li, Z., Huang, W., Yuan, J., et al. (2012). SOAPdenovo2: an empirically improved memory-efficient short-read de novo assembler. Gigascience 1(1), 18. doi: 10.1186/2047-217X-1-18.

Lye, H.-S., Rahmat-Ali, G.R., and Liong, M.-T. (2010a). Mechanisms of cholesterol removal by lactobacilli under conditions that mimic the human gastrointestinal tract. International Dairy Journal 20(3), 169–175. doi: 10.1016/j.idairyj.2009.10.003.

Lye, H.S., Rusul, G., and Liong, M.T. (2010b). Removal of cholesterol by lactobacilli via incorporation and conversion to coprostanol. J Dairy Sci 93(4), 1383–1392. doi: 10.3168/jds.2009-2574.

M, S. (1990). Identification of bacteria by gas chromatography of cellular fatty acids, MIDI technical note 101. MIDI Inc, Newark, DE.

Mercan, E., Ispirli, H., Sert, D., Yilmaz, M.T., and Dertli, E. (2015). Impact of exopolysaccharide production on functional properties of some *Lactobacillus salivarius* strains. Arch Microbiol 197(9), 1041–1049. doi: 10.1007/s00203-015-1141-0.

Morotomi, M., Nagai, F., and Watanabe, Y. (2012). Description of *Christensenella minuta* gen. nov., sp. nov., isolated from human faeces, which forms a distinct branch in the order *Clostridiales*, and proposal of *Christensenellaceae* fam. nov. Int J Syst Evol Microbiol 62(Pt 1), 144–149. doi: 10.1099/ijs.0.026989-0.

Ndongo, S., Dubourg, G., Khelaifia, S., Fournier, P.E., and Raoult, D. (2016a). *Christensenella timonensis*, a new bacterial species isolated from the human gut. New Microbes New Infect 13, 32–33. doi: 10.1016/j.nmni.2016.05.010.

Ndongo, S., Khelaifia, S., Fournier, P.E., and Raoult, D. (2016b). *Christensenella massiliensis*, a new bacterial species isolated from the human gut. New Microbes New Infect 12, 69–70. doi: 10.1016/j.nmni.2016.04.014.

Palmer, C., Bik, E.M., DiGiulio, D.B., Relman, D.A., and Brown, P.O. (2007). Development of the human infant intestinal microbiota. PLoS Biol 5(7), e177. doi: 10.1371/journal.pbio.0050177.

Pan, D.D., Zeng, X.Q., and Yan, Y.T. (2010). Characterisation of *Lactobacillus fermentum* SM-7 isolated from koumiss, a potential probiotic bacterium with cholesterol-lowering effects. J Sci Food Agric 91(3), 512–518. doi: 10.1002/jsfa.4214.

Pineiro, M., and Stanton, C. (2007). Probiotic bacteria: legislative framework--requirements to evidence basis. J Nutr 137(3 Suppl 2), 850S–853S.

Rosa, B.A., Hallsworth-Pepin, K., Martin, J., Wollam, A., and Mitreva, M. (2017). Genome Sequence of *Christensenella minuta* DSM 22607^T^. Genome Announc 5(2). doi: 10.1128/genomeA.01451-16.

Rudel, L.L., Morris, M. D. (1973). Determination of cholesterol using O-phthalaldehyde. J Lipid Res 14(3), 364–366.

Rzhetsky, A., and Nei, M. (1993). Theoretical foundation of the minimum-evolution method of phylogenetic inference. Mol Biol Evol 10(5), 1073–1095.

Saitou, N., and Nei, M. (1987). The neighbor-joining method: a new method for reconstructing phylogenetic trees. Mol Biol Evol 4(4), 406–425.

Smibert RM, K.N. (1994). Phenotypic Characterization. Methods for General and Molecular Bacteriology: American Society for Microbiology.

Sorokin, D.Y. (2005). Is there a limit for high-pH life? Int J Syst Evol Microbiol 55(Pt 4), 1405–1406. doi: 10.1099/ijs.0.63737-0.

Tamura, K., Peterson, D., Peterson, N., Stecher, G., Nei, M., and Kumar, S. (2011). MEGA5: molecular evolutionary genetics analysis using maximum likelihood, evolutionary distance, and maximum parsimony methods. Mol Biol Evol 28(10), 2731–2739. doi: 10.1093/molbev/msr121.

Thompson, J.D., Higgins, D.G., and Gibson, T.J. (1994). CLUSTAL W: improving the sensitivity of progressive multiple sequence alignment through sequence weighting, position-specific gap penalties and weight matrix choice. Nucleic Acids Res 22(22), 4673–4680.

Tindall, B.J., Rossello-Mora, R., Busse, H.J., Ludwig, W., and Kampfer, P. (2010). Notes on the characterization of prokaryote strains for taxonomic purposes. Int J Syst Evol Microbiol 60(Pt 1), 249–266. doi: 10.1099/ijs.0.016949-0.

Tittsler RP, S.L. (1936). The Use of Semi-solid Agar for the Detection of Bacterial Motility. J Bacteriol. 31(6), 6.

Tok, E., and Aslim, B. (2010). Cholesterol removal by some lactic acid bacteria that can be used as probiotic. Microbiology and Immunology. doi: 10.1111/j.1348-0421.2010.00219.x.

Tremaroli, V., and Backhed, F. (2012). Functional interactions between the gut microbiota and host metabolism. Nature 489(7415), 242–249. doi: 10.1038/nature11552.

Tsai, C.C., Lin, P.P., Hsieh, Y.M., Zhang, Z.Y., Wu, H.C., and Huang, C.C. (2014). Cholesterol-lowering potentials of lactic acid bacteria based on bile-salt hydrolase activity and effect of potent strains on cholesterol metabolism in vitro and in vivo. ScientificWorldJournal 2014, 690752. doi: 10.1155/2014/690752.

Turnbaugh, P.J., Backhed, F., Fulton, L., and Gordon, J.I. (2008). Diet-induced obesity is linked to marked but reversible alterations in the mouse distal gut microbiome. Cell Host Microbe 3(4), 213–223. doi: 10.1016/j.chom.2008.02.015.

Turnbaugh, P.J., Ley, R.E., Mahowald, M.A., Magrini, V., Mardis, E.R., and Gordon, J.I. (2006). An obesity-associated gut microbiome with increased capacity for energy harvest. Nature 444(7122), 1027–1031. doi: 10.1038/nature05414.

Turnbaugh, P.J., Ridaura, V.K., Faith, J.J., Rey, F.E., Knight, R., and Gordon, J.I. (2009). The effect of diet on the human gut microbiome: a metagenomic analysis in humanized gnotobiotic mice. Sci Transl Med 1(6), 6ra14. doi: 10.1126/scitranslmed.3000322.

Ventura, M. (2009). Microbial diversity in the human intestine and novel insights from metagenomics. Frontiers in Bioscience Volume(14), 3214. doi: 10.2741/3445.

Welman, A.D., and Maddox, I.S. (2003). Exopolysaccharides from lactic acid bacteria: perspectives and challenges. Trends Biotechnol 21(6), 269–274. doi: 10.1016/S0167-7799(03)00107-0.

Yoon, H.S., Ju, J.H., Kim, H.N., Park, H.J., Ji, Y., Lee, J.E., et al. (2013). Reduction in cholesterol absorption in Caco-2 cells through the down-regulation of Niemann-Pick C1-like 1 by the putative probiotic strains *Lactobacillus rhamnosus* BFE5264 and *Lactobacillus plantarum* NR74 from fermented foods. Int J Food Sci Nutr 64(1), 44–52. doi: 10.3109/09637486.2012.706598.

Yoon, S.H., Ha, S.M., Kwon, S., Lim, J., Kim, Y., Seo, H., et al. (2017). Introducing EzBioCloud: a taxonomically united database of 16S rRNA gene sequences and whole-genome assemblies. Int J Syst Evol Microbiol 67(5), 1613–1617. doi: 10.1099/ijsem.0.001755.

Zou, Y., Liu, F., Fang, C., Wan, D., Yang, R., Su, Q., et al. (2013). *Lactobacillus shenzhenensis* sp. nov., isolated from a fermented dairy beverage. Int J Syst Evol Microbiol 63(Pt 5), 1817–1823. doi: 10.1099/ijs.0.041111-0.

Zou, Y., Xue, W., Luo, G., Deng, Z., Qin, P., Guo, R., et al. (2019). 1,520 reference genomes from cultivated human gut bacteria enable functional microbiome analyses. Nat Biotechnol 37(2), 179–185. doi: 10.1038/s41587-018-0008-8.

