## Supplementary figures and images for "Taxonomic description and genome sequence of *Christensenella intestinihominis* sp. nov., a novel cholesterol-lowering bacterium isolated from human gut"

### Supplementary Figure S1

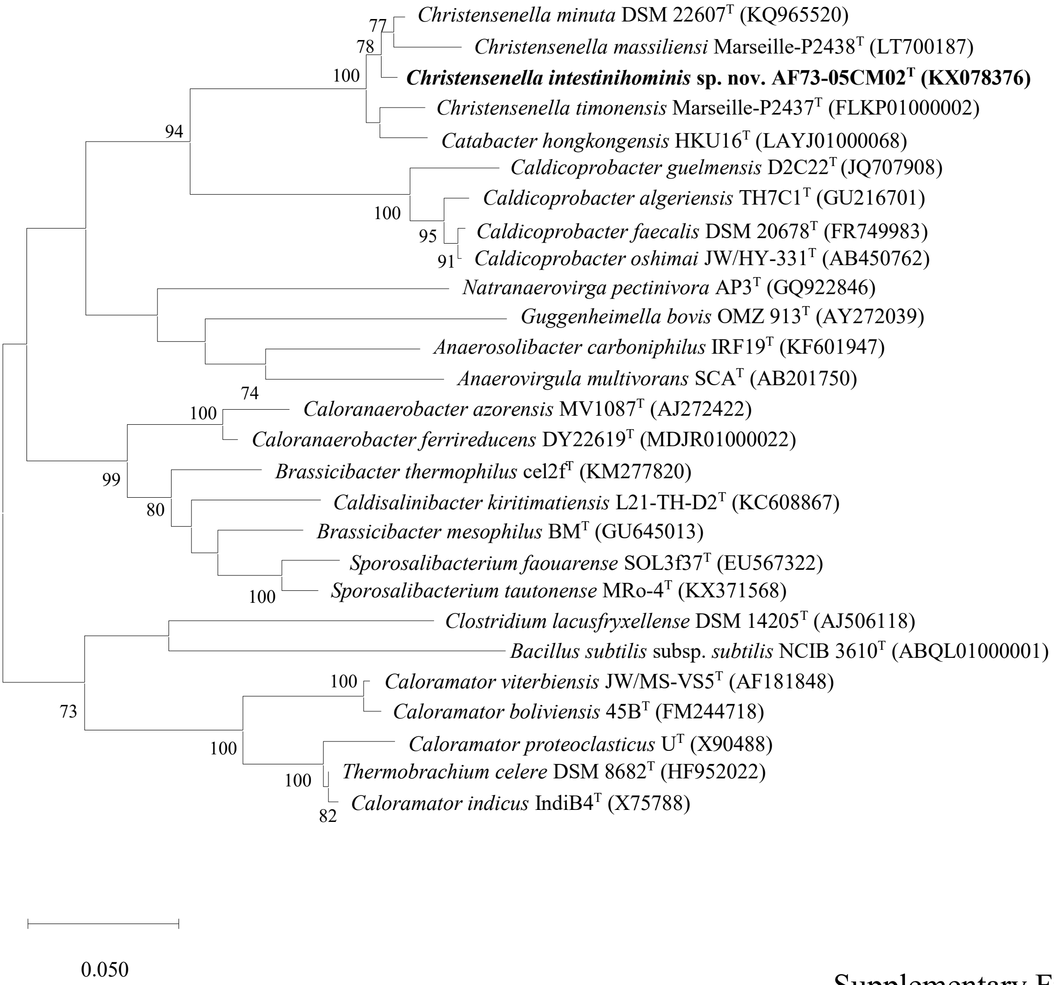


Supplementary Figure S1

### Supplementary Figure S2

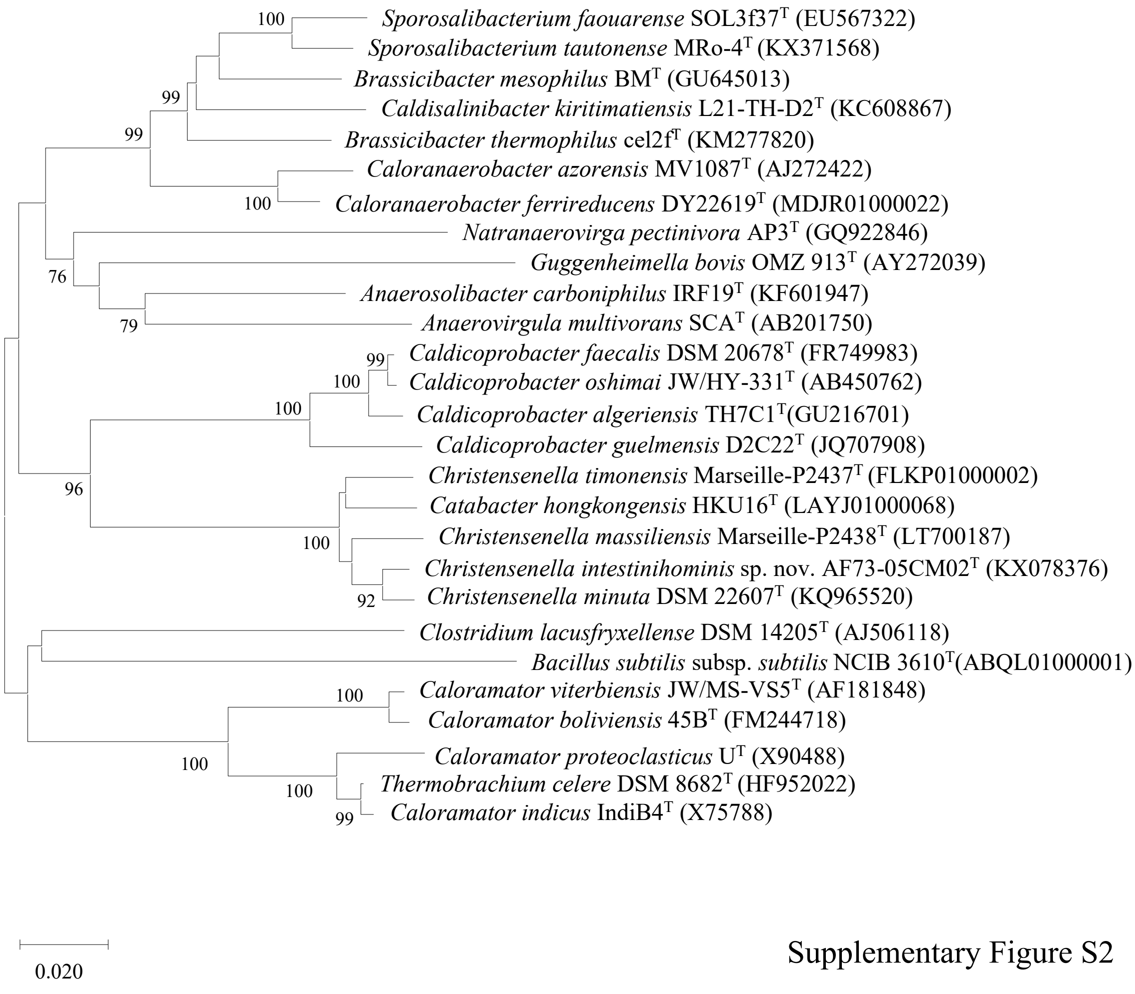


Supplementary Figure S2
