## Supplementary Table S1 for "Taxonomic description and genome sequence of *Christensenella intestinihominis* sp. nov., a novel cholesterol-lowering bacterium isolated from human gut"

**Supplementary Table S1. Number of genes associated with general COG functional categories in the** **genome of *C. intestinihominis* AF73-05CM02^T^ and *C. minuta* DSM 22607^T^.**

| **COG Categories** | **Code** | **AF73-05CM02^T^** | | **DSM 22607^T^** | |
| --- | --- | --- | --- | --- | --- |
|  |  | **Gene Count** | **% of Total Genes** | **Gene Count** | **% of Total Genes** |
| Amino acid transport and metabolism | **E** | 159 | 6.0 | 164 | 6.3 |
| Carbohydrate transport and metabolism | **G** | 286 | 10.8 | 283 | 10.8 |
| Cell cycle control, cell division, chromosome partitioning | **D** | 33 | 1.2 | 31 | 1.2 |
| Cell motility | **N** | 49 | 1.9 | 18 | 0.7 |
| Cell wall/membrane/envelope biogenesis | **M** | 106 | 4.0 | 106 | 4.1 |
| Coenzyme transport and metabolism | **H** | 67 | 2.5 | 70 | 2.7 |
| Cytoskeleton | **Z** | 1 | 0.04 | 0 | 0 |
| Defense mechanisms | **V** | 57 | 2.2 | 60 | 2.3 |
| Energy production and conversion | **C** | 148 | 5.6 | 141 | 5.4 |
| Extracellular structures | **W** | 7 | 0.3 | 7 | 0.3 |
| Function unknown | **S** | 88 | 3.3 | 90 | 3.4 |
| General function prediction only | **R** | 176 | 6.7 | 182 | 7.0 |
| Inorganic ion transport and metabolism | **P** | 76 | 2.9 | 70 | 2.7 |
| Intracellular trafficking, secretion, and vesicular transport | **U** | 32 | 1.2 | 29 | 1.1 |
| Lipid transport and metabolism | **I** | 65 | 2.5 | 87 | 3.3 |
| Mobilome: prophages, transposons | **X** | 17 | 0.6 | 20 | 0.8 |
| Nucleotide transport and metabolism | **F** | 68 | 2.6 | 67 | 2.6 |
| Posttranslational modification, protein turnover, chaperones | **O** | 67 | 2.5 | 64 | 2.5 |
| Replication, recombination and repair | **L** | 98 | 3.7 | 102 | 3.9 |
| Secondary metabolites biosynthesis, transport and catabolism | **Q** | 44 | 1.7 | 56 | 2.1 |
| Signal transduction mechanisms | **T** | 151 | 5.7 | 122 | 4.7 |
| Transcription | **K** | 197 | 7.5 | 187 | 7.2 |
| Translation, ribosomal structure and biogenesis | **J** | 184 | 7.0 | 175 | 6.7 |
| Not in COGs |  | 466 | 17.6 | 478 | 18.3 |
