## Supplementary Table S3 for "Taxonomic description and genome sequence of *Christensenella intestinihominis* sp. nov., a novel cholesterol-lowering bacterium isolated from human gut"

**Supplementary Table S3.** **Enzymatic characteristics of strain AF73-05CM02^T^ from API ZYM test.**

| **API ZYM** | **AF73-05CM02^T^** |
| --- | --- |
| Alkaline phosphatase | – |
| Esterase(C4) | – |
| Esterase lipase(C8) | – |
| Lipase (C14) | – |
| Leucine arylamidase | – |
| Valine arylamidase | – |
| Cystine arylamidase | – |
| Trypsin | – |
| α-chymotrypsin | – |
| Acid phosphatase | – |
| Naphthol-AS-B1-phosphohydrolase | + |
| α-galactosidase | – |
| β-galactosidase | – |
| β- glucuronidase | – |
| α-glucosidase | – |
| β- glucosidase | – |
| N-acetyl-β- glucosaminidase | – |
| α-mannosidase | – |
| α-fucosidase | – |

+, Positive; –, negative.
