## Supplementary Table S4 for "Taxonomic description and genome sequence of *Christensenella intestinihominis* sp. nov., a novel cholesterol-lowering bacterium isolated from human gut"

**Supplementary Table S4. Carbon source assimilation of strain AF73-05CM02^T^ from** **API 20A and API 50CHL test.**

| **Carbon substrate (API 20A)** | **AF73-05CM02^T^** | **Carbon substrate (API 20A)** | **AF73-05CM02^T^** |
| --- | --- | --- | --- |
| d-Glucose | + | Glycerol | − |
| d-Mannose | w | d-Cellobiose | − |
| d-Lactose | − | d-Mannitol | − |
| d-Sucrose | + | d-Melezitose | w |
| d-Maltose | w | d-Raffinose | w |
| Salicin | w | d-Sorbitol | + |
| d-Xylose | + | l-Rhamnose | + |
| l-Arabinose | + | d-Trehalose | − |

| **Carbon substrate (API 50CHL)** | **AF73-05CM02^T^** | **Carbon substrate (API 50CHL)** | **AF73-05CM02^T^** |
| --- | --- | --- | --- |
| Glycerol | − | Salicin | + |
| Erythritol | w | Cellobiose | − |
| d-Arabinose | + | d-Maltose | w |
| l-Arabinose | + | d-Lactose | − |
| d-Ribose | + | d-Melibiose | − |
| d-Xylose | + | d-Sucrose | + |
| l-Xylose | w | d-Trehalose | − |
| d-Adonitol | − | Inulin | − |
| Methyl-*β*-d-Xylopyranoside | + | d-Melezitose | w |
| d-Galactose | + | d-Raffinose | w |
| d-Glucose | + | Starch | − |
| d-Fructose | + | Glycogen | − |
| d-Mannose | + | Xylitol | + |
| l-Sorbose | + | Gentiobiose | + |
| l-Rhamnose | + | d-Turanose | + |
| Dulcitol | − | d-Lyxose | + |
| Inositol | − | d-Tagatose | + |
| d-Mannitol | − | d-Fucose | + |
| d-Sorbitol | + | l-Fucose | − |
| Methyl-α-D-Mannopyranoside | − | d-Arabitol | − |
| Methyl-d-Glucopyranoside | − | l-Arabitol | − |
| N-acetyl-Glucosamine | − | Gluconate | − |
| Amygdalin | − | 2-Ketogluconate | − |
| Arbutin | + | 5-Ketogluconate | − |

+, positive; −, negative; w, weakly positive reaction.
