## Supplementary Table S5 for "Taxonomic description and genome sequence of *Christensenella intestinihominis* sp. nov., a novel cholesterol-lowering bacterium isolated from human gut"

**Supplementary Table S5. Antibiotic sensitivity of strain AF73-05CM02^T^.**

| **NO.** | **Antibiotic** | **Concentration (μg)** | **AF73-05CM02^T^** |
| --- | --- | --- | --- |
| 1 | Penicillin | 10 | S |
| 2 | Oxacillin | 1 | R |
| 3 | Ampicillin | 10 | S |
| 4 | Carbenicillin | 100 | S |
| 5 | Piperacillin | 100 | S |
| 6 | Vancomycin | 30 | S |
| 7 | Polymyxin B | 300 IU | S |
| 8 | Compound sulfamethoxazole | 25 | R |
| 9 | Furazolidone | 300 | S |
| 10 | Chloroamphenicol | 30 | S |
| 11 | Clindamycin | 2 | S |

R, resistant; S, sensitive.
