## Supplementary Table S6 for "Taxonomic description and genome sequence of *Christensenella intestinihominis* sp. nov., a novel cholesterol-lowering bacterium isolated from human gut"

**Supplementary Table S6. Metabolic end products from glucose for strain AF73-05CM02^T^ and *C. minuta* DSM 22607^T^.**

| **Metabolic end products (mmol/L)** | | **AF73-05CM02^T^** | **DSM 22607** |
| --- | --- | --- | --- |
| **Short-chain fatty acids (SCFAs)** | Formic acid | 5.21 | 4.64 |
|  | Acetic acid | 14.60 | 33.97 |
|  | Propionic acid | 0 | 0 |
|  | Isobutyric acid | 0.37 | 0.37 |
|  | Butyric acid | 1.63 | 6.65 |
|  | Valeric acid | 0.36 | 0.36 |
|  | Isovaleric acid | 0 | 0 |
| **Organic acids** | Quinic acid | 0 | 0 |
|  | L-(+) Lactic acid | 4.13 | 7.83 |
|  | Oxalic acid | 0 | 0 |
|  | Malonic acid | 0 | 0.08 |
|  | Benzoic acid | 0 | 5.00 |
|  | Maleic acid | 0 | 0 |
|  | Fumaric acid | 0 | 0 |
|  | D-(+)-Malic acid | 0 | 0 |
|  | Adipic acid | 0.24 | 0.28 |
|  | L-(-)-Tartaric acid | 0 | 0 |
|  | Shikimic acid | 0 | 0 |
|  | L-Ascorbic acid | 0.11 | 0.11 |
|  | Citric acid | 0.18 | 0.14 |
|  | DL-Isocitric acid trisodium salt hydrate | 0 | 0 |
